# Research on the Estimation Model of Seedling Sowing Period Based on Different Altitude

**DOI:** 10.64898/2025.11.29.691278

**Authors:** Yabo Jin, Zhenguo Wang, Jianqin Luo

## Abstract

The selection of sowing date is a key link in tobacco planting, and an appropriate sowing date can affect the survival rate and agronomic traits of tobacco plants. This study collected data on tobacco seedlings during the sowing and transplanting periods in the southwestern region of China, combined with altitude gradient data of the planting area. Time series analysis was used to construct a tobacco seedling sowing period estimation model and establish an application platform. The model was validated by comparing the survival rate of transplanting. The results indicate that the proposed LSTM model has an R2 of 0.893, a MAE of 1.796, an RSME of 2.221, and a validation model accuracy of 93.3%. The research results provide scientific sowing time basis for tobacco growers in different altitude regions, and results provide scientific sowing time basis for tobacco growers in different altitude regions, and offer suggestions for optimizing tobacco planting management and promoting the sustainable development of the tobacco industry.

## 1. Introduction

As a vital cash crop, tobacco plays a significant role in global agriculture and trade. This annual herb of the Solanaceae family (Tobacco genus) has unique biological traits and growth requirements. Research indicates that tobacco seedlings thrive best at temperatures between 18°C and 25°C, with extreme heat or cold severely impacting their growth rate and quality ^[1]^. Adequate sunlight is essential, requiring at least 8 hours of daily exposure. Altitude affects temperature, light intensity, and soil conditions, which in turn influence the tobacco growth cycle and quality. In high-altitude regions, cooler temperatures and stronger sunlight may extend the growth cycle while potentially enhancing leaf quality ^[2]^.

Currently, seedling growers address the impact of altitude gradients on tobacco seedling growth by controlling cultivation duration. Building on this approach, this study employs Long Short-Term Memory (LSTM) networks to develop a tobacco seedling sowing period prediction model. As a specialized recurrent neural network, LSTM effectively resolves gradient vanishing and explosion issues in traditional RNNs when processing long sequences ^[3]^. By incorporating memory units and gating mechanisms, LSTM captures temporal dependencies in extended data streams, making it suitable for time series prediction tasks. This model has been widely applied in forecasting fields ^[4]^. For instance, Sun Borui proposed an LSTM-based crop water requirement prediction model that uses meteorological data (air temperature/humidity, wind speed, sunshine duration) as input features and crop water demand as output labels, significantly improving prediction accuracy and providing scientific basis for agricultural irrigation management ^[5]^. Researchers have also developed an LSTM rice yield prediction model using improved principal component analysis (PCA), which reduces input data dimensionality to eliminate noise and redundancy while retaining key information, thereby enhancing yield prediction precision. The application of LSTM models in crop soil moisture demand prediction has achieved notable results ^[6]^. For example, a soil moisture dynamic model combining Transformer and LSTM algorithms accurately predicts soil moisture variations in the Hetao Irrigation District, offering scientific support for rational agricultural water resource planning and utilization ^[7]^.

The duration of tobacco seedling cultivation is influenced by multiple factors across different altitude gradients, including temperature, humidity, and light exposure. These data are typically presented in time series format ^[8]^. By analyzing historical patterns, the LSTM model can deduce the optimal sowing period for tobacco seedlings based on their growth conditions, thereby providing production guidance. The research model has been developed into a digital application platform that inputs the transplanting time of mature seedlings and altitude gradients to output estimated sowing times ^[9]^. This system holds significant implications for optimizing tobacco cultivation management.

## 2 Materials and Methods

### 2.1 Setting of the Research Area

The tobacco seedling trials were conducted in three key regions: Fengjie Tobacco Area in northeastern Chongqing, Baise Tobacco Area in Guangxi, and Nanxiong Tobacco Area in Guangdong. These areas feature diverse terrains including plains, hills, and mountains with varying elevations. The primary tobacco cultivar used was “Yunyan 87”. In the northeastern Chongqing region, three representative plains and hilly areas were selected as trial sites, each containing three replicate plots measuring 666 square meters. The Qinba Mountainous Area in this region had elevations ranging from 495 to 1,570 meters. The Baise Tobacco Area in Guangxi, predominantly hilly and mountainous, featured six trial sites with three replicate plots per site, each covering 100 square meters. The hilly areas had elevations between 350 and 800 meters, while the mountainous areas spanned 600 to 1,100 meters. The Nanxiong Tobacco Area in Guangdong encompassed both plains and hills, with six trial sites each containing three replicate plots of 100 square meters. The plains had elevations ranging from 180 to 280 meters, while the hilly areas extended from 280 to 650 meters.

### 2.2 Sowing and Transplanting

The sowing schedule was preliminarily determined based on local climate conditions and traditional cultivation practices, with multiple sowing date treatments established at different experimental sites. Each treatment was spaced 5 days apart, resulting in a total of 5 sowing date treatments. Manual spot sowing was employed. Transplanting timing was determined according to the growth status of tobacco seedlings and local climate conditions, selecting healthy seedlings free from pests and diseases. The transplanting density was set at 18,000 plants per hectare. After transplantation, timely irrigation for root establishment was conducted, followed by appropriate shading to enhance seedling survival rates.

### 2.3 Data Collection

For each experimental plot, meticulously record the sowing date, transplanting date, and growth status of tobacco seedlings at transplanting, including key agronomic traits such as height, stem diameter, and leaf count. Conduct survival rate surveys 30-40 days post-transplantation, then calculate the survival rate using the formula: Survival Rate (%) = (Number of Surviving Seedlings / Total Transplanted Seedlings) × 100. Employ high-precision GPS devices to measure the altitude of each plot for data accuracy. Additionally, document the geographical coordinates of each plot to facilitate subsequent analysis of altitude-related variations.

### 2.4 Data Processing and Model Construction

The collected data on sowing period, transplanting period, transplanting survival rate, and altitude were organized to establish a database. Preliminary analysis was conducted, including descriptive statistical analysis and calculation of average values for each indicator to understand the basic characteristics of the data. Logistic regression analysis was performed to examine the relationship between sowing period and altitude. This preliminary assessment of their correlation provided a basis for subsequent model construction.

The sowing and transplanting period data were organized chronologically to form a time series dataset. Using time series analysis methods, the dataset was examined for stationarity and seasonality. Based on the analysis results, an LSTM model was constructed. This model uses altitude, sowing period, and planned transplanting period as input variables, with sowing period prediction as the output variable, forming a multi-input single-output LSTM architecture ^[10]^. Through training, the model learns the complex nonlinear relationships between sowing/transplanting periods and transplanting survival rates across different altitudes.

The preprocessed dataset was divided into a training set (70%) and a test set (30%). The LSTM model was trained using the training set, and the model was fine-tuned with the validation set to select optimal model parameters ^[11]^.

### 2.5 Model Verification and Platform Development

The model’s predictive performance was evaluated using the test set, with metrics such as R ^2^, MAE, and RMSE calculated to assess its accuracy and stability ^[12]^. Additionally, the model’s predicted sowing dates were compared with actual transplant survival rates to verify its accuracy, further validating the model’s practicality.

The model is implemented as a system application built on the J2EE platform, a distributed application environment that deploys computing systems as distributed components across multiple servers ^[13]^. The system integrates the Spring framework with a front-end and back-end separation architecture. It incorporates REST API open interfaces ^[14]^, IDBC database applications ^[15]^, IotDB for time-series data storage ^[16]^, and Jon document model configurations ^[17]^.

## 3 Results

### 3.1 Agronomic traits and survival rate of tobacco plants

The study collected 700 valid data sets from three planting bases. The highest altitude in the study area was 1,350 meters, and the lowest was 242 meters. In all surveyed plots, the average height of tobacco plants (measured from the ridge furrows) was 84.61 ± 13.52cm, the stem diameter was 2.45 ± 0.37cm, the number of effective leaves was 16.52 ± 2.71, the maximum leaf length was 37.58±6.1cm, the maximum leaf width was 23.49±3.8cm, and the survival rate was 91.93±5.89%. The agronomic traits of the tobacco plants met normal standards, and the survival rate met production requirements. Regression analysis was conducted between the agronomic traits of tobacco plants, the survival rate, and different altitude gradients to analyze their effects on tobacco plants at different altitudes. The results showed (Figure 1) that among the agronomic traits of tobacco plants, plant height had a certain inverse relationship with altitude gradient, generally indicating that the higher the altitude, the relatively lower the plant height, R^2^=0.24, suggesting that this correlation was not strong. The stem diameter showed no linear relationship with altitude gradient, R ^2^ only 0.12, while the number of effective leaves had a certain inverse relationship with altitude gradient (R^2^=0.39), meaning the higher the altitude gradient, the relatively fewer the effective leaves. The analysis results also indicated that the maximum leaf length and width of tobacco plants were significantly associated with altitude. The survival rate of tobacco plants had a certain inverse relationship with altitude gradient (R^2^=0.34), meaning the higher the altitude, the relatively lower the survival rate.

**Figure 1.**
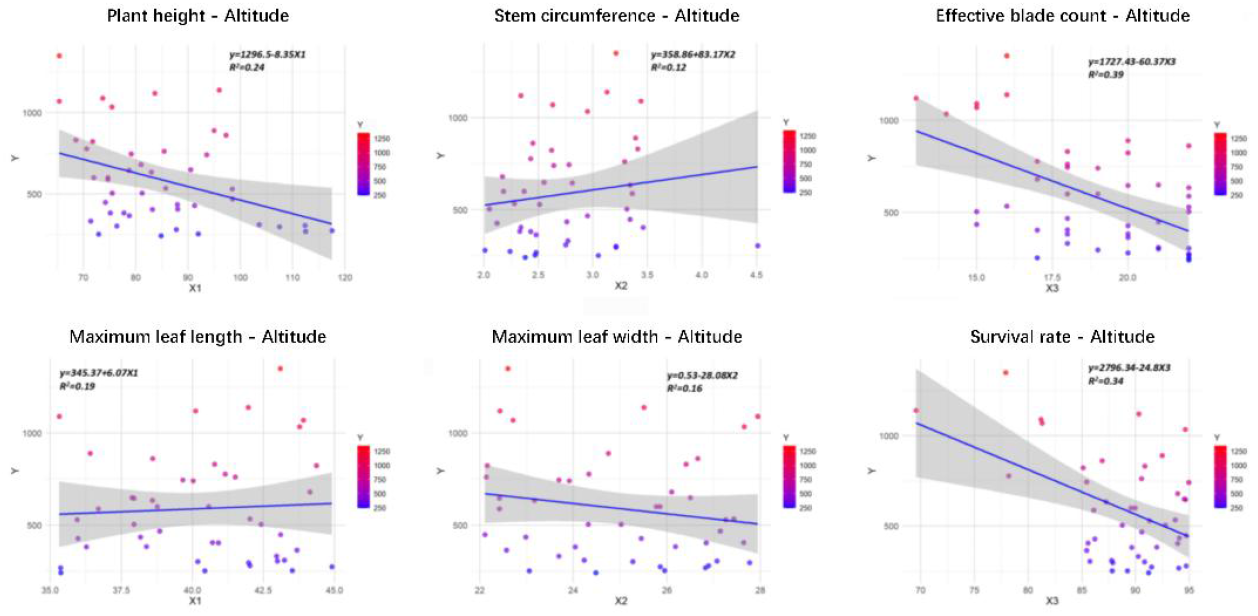
Results of agronomic traits and survival rate of tobacco plants at different altitude gradients

### 3.2 Establishment of Long and Short-Term Memory Network Model

During model training, 1,500 iterations were performed. Through parameter tuning and optimization, the final optimal model parameters and their selection criteria are presented in Table 1. The Long Short-Term Memory (LSTM) network model’s performance in establishing tobacco plant transplanting time models at different altitudes (Figure 2) shows that predicted values generally align closely with actual values across most samples, though some exhibit significant deviations ^[18]^. The Root Mean Square Error (RMSE) of 2.6128 indicates the average prediction error. Similar to the training dataset, the RMSE for the test dataset is 2.2217, reflecting minimal deviations in certain cases. Overall, both training and test datasets demonstrate high consistency between predicted and actual values, indicating low prediction errors.

**Table 1.**
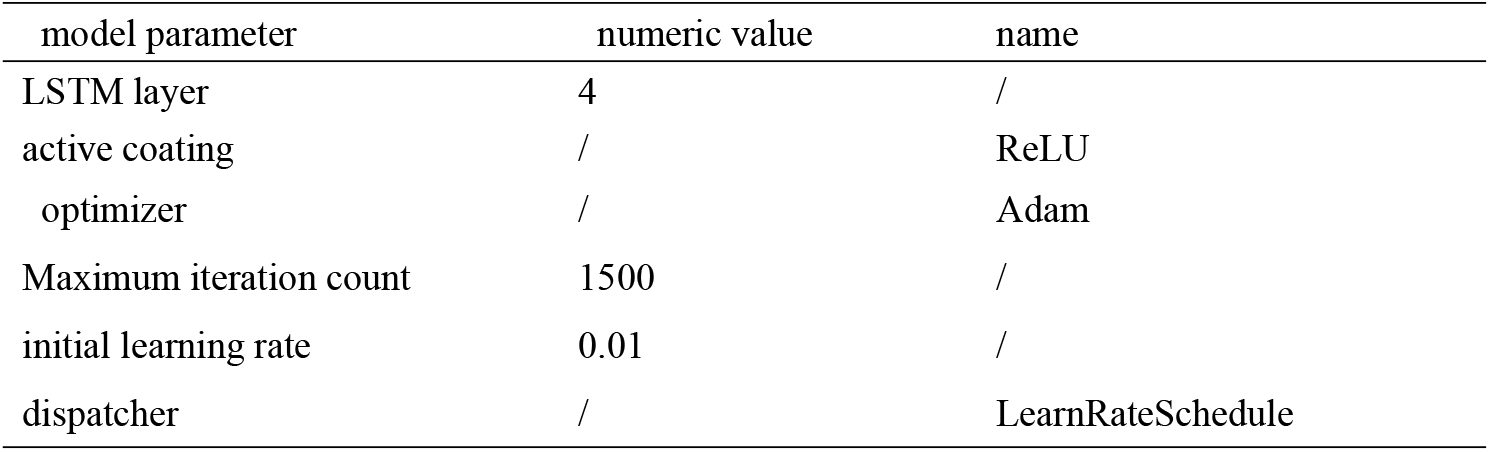
Optimal parameter values of long and short-term memory network model.

**Figure 2.**
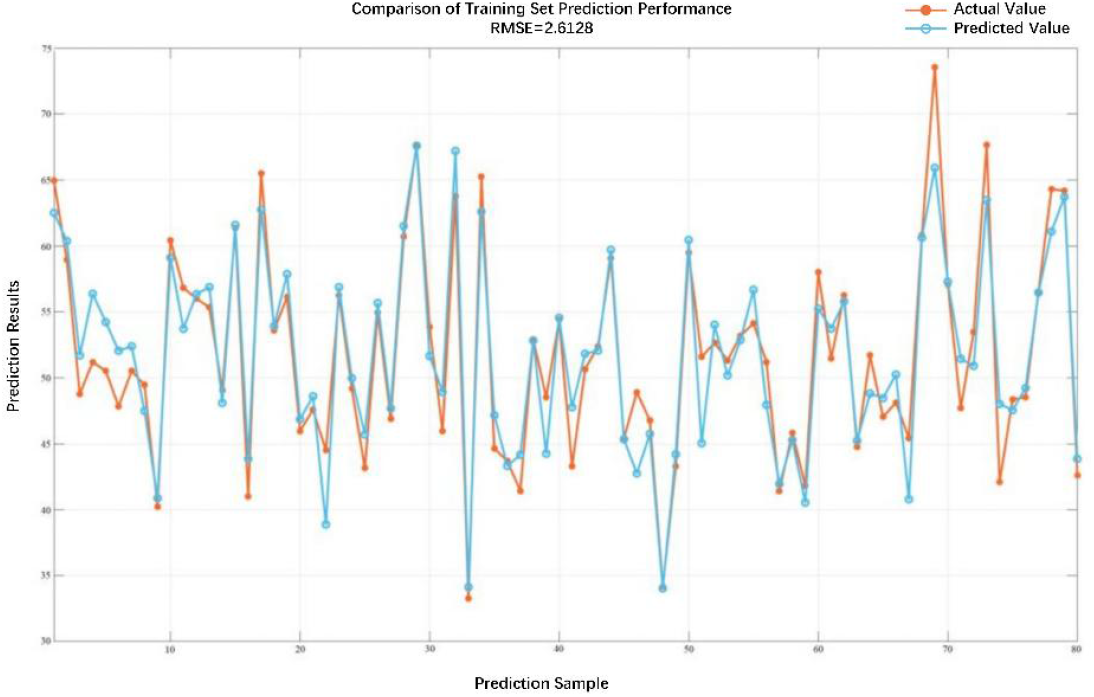
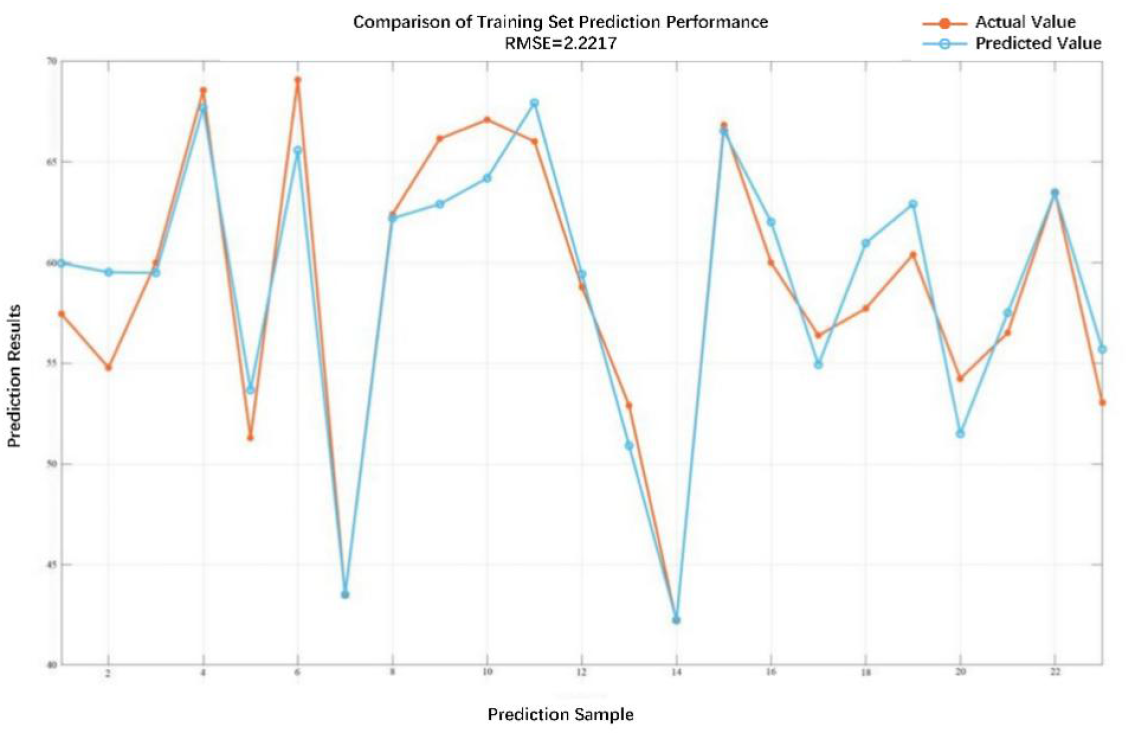
Results of Long-Term and Short-Term Memory Network Model for Sowing Period at Different Altitude Gradients (X-axis: randomly selected prediction and training samples; Y-axis: time from sowing to field transplanting of tobacco seedlings at different altitudes)

The comparison of fitting residuals between the model’s true values and predicted values (Figure 3) presents a scatter plot of predicted values (y-axis) and true values (x-axis) from the training set. In an ideal scenario, all points should align perfectly along the diagonal (dashed line), indicating perfect consistency between predictions and actual measurements. The graph shows most points clustered near the diagonal with only a few significant deviations, suggesting some prediction errors in the training set. Similarly, the test set data reveals most points concentrated near the diagonal with minor outliers, indicating residual prediction errors. The Root Mean Square Error (RMSE) value of 2.2217 for the test set demonstrates better generalization performance (2.2217 vs. 2.6128) compared to the training set, indicating the model’s ability to extrapolate to unseen data. The model achieved an R^2^ coefficient of 0.865 on the training set and 0.893 on the test set, with Mean Absolute Error (MAE) values of 1.963 and 1.796 respectively. The LSTM model consistently performed well in predicting sowing periods across different altitudes, showing minimal discrepancies between predicted and true values. Validation using seedling time models at varying altitude gradients achieved an accuracy rate of 93.35±2.54%, demonstrating excellent predictive performance.

**Figure 3.**
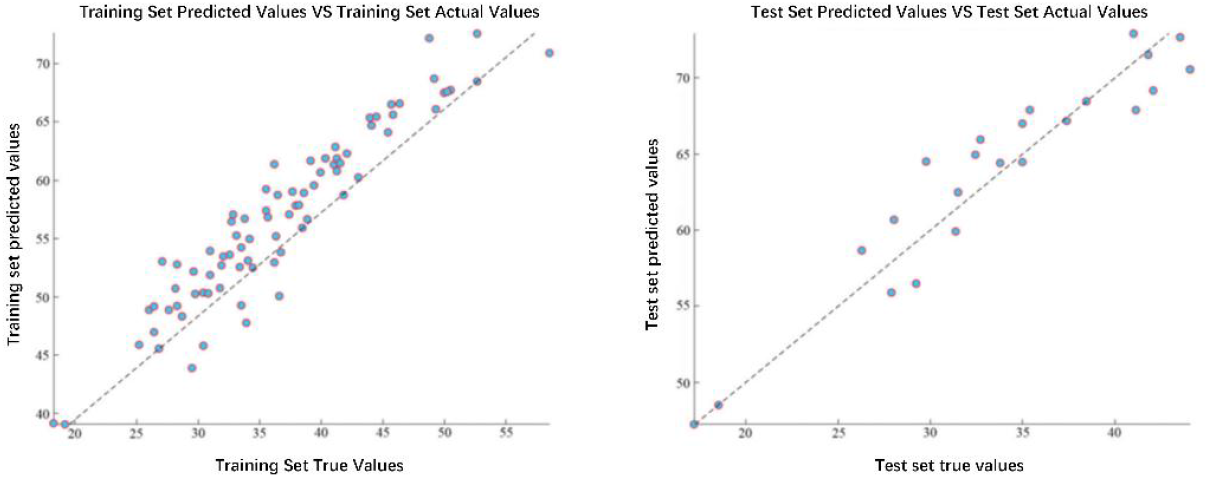
Comparison of real values and predicted values between the training set and test set in the long short-term memory network model

### 3.3 Application Platform Deployment

The model’s platform deployment was optimized for real-world production, with the planned sowing time now formatted as **year**/**month**/**day** (revised as Planned Transplantation Date) for easier selection during operations. The UI development, shown in Figure 4, includes the model engine configuration parameters and computational logic.

**Figure 4.**
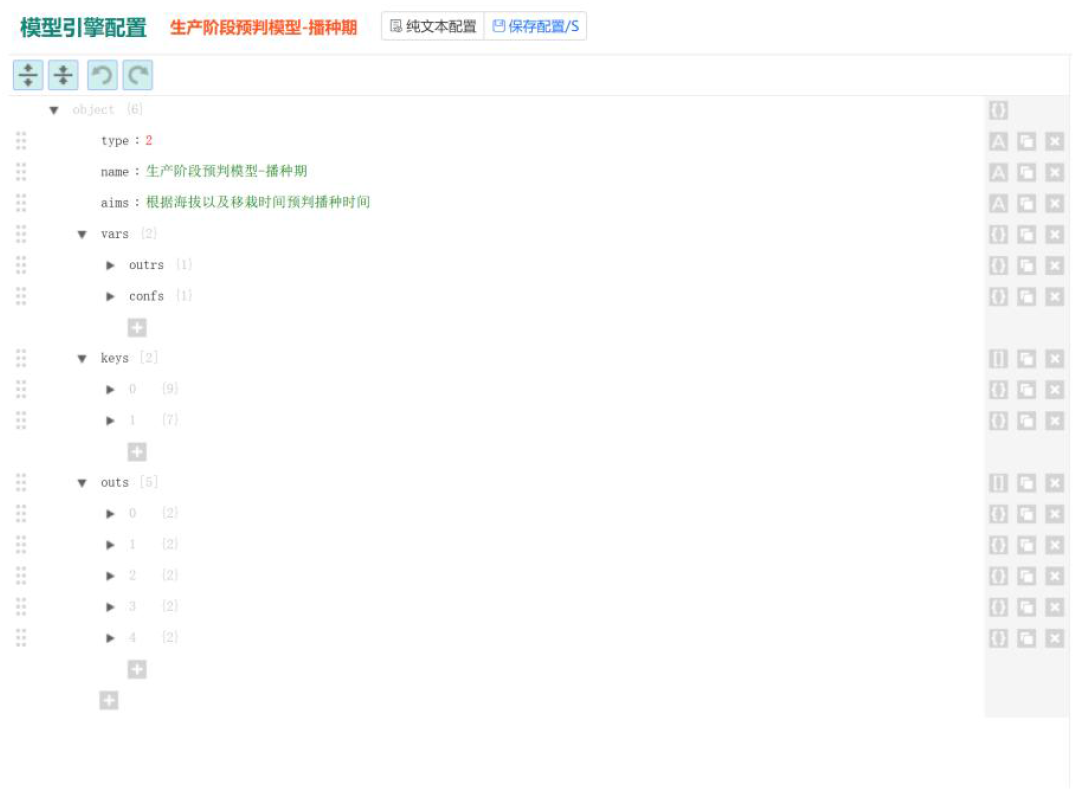
Long Short-Term Memory Network Model Based on Cloud Platform Deployment Logic

The deployed model was tested, with input and output results shown in Figure 5. The model’s input interface on the left contains data collection dates, bases, and stations. For this test, the collection base was selected as the Chongqing Fengjie Tobacco Planting Base near Dashibao Tobacco Station. The altitude of this area was measured by the 484GPS module during IoT device connectivity testing, recording 762 meters ^[19]^. The simulation simulated tobacco seedling transplanting operations in the field on May 1, 2025. After inputting the numerical data, the model’s computational output appeared on the right. Based on the model’s calculations, the results were adjusted by adding 2 days before and after the original date (typically 5 days for sowing operations), resulting in a sowing period from March 12 to March 17,2025. The upper-right section displays the model’s call history records, enabling operators to generate statistical reports for sowing coverage within their jurisdiction.

**Figure 5.**
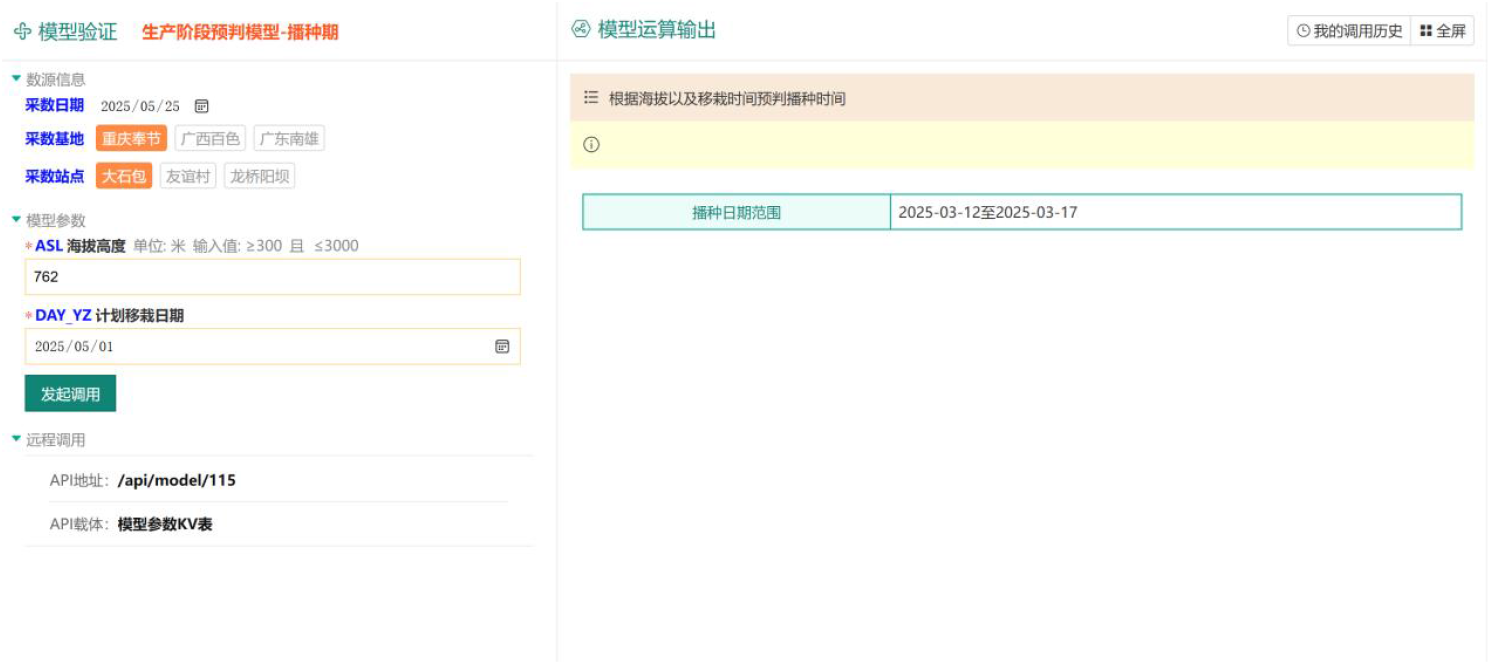
Interface of the model application system for seedling sowing period

## 4 Discussion

At present, tobacco researches mainly focus on the prediction of transplanting period, but few methods can predict the sowing period directly. Therefore, it is of great significance to develop a more accurate model to predict the sowing period for improving the efficiency and quality of tobacco production ^[20]^.

The tobacco plant sowing period prediction model assists farmers in determining optimal planting schedules to ensure proper growth under suitable environmental conditions. As altitude significantly impacts crop development — particularly in southern tobacco-growing regions dominated by mountainous terrain—it is crucial to develop sowing period prediction models for different altitude gradients ^[21]^. Temperature generally decreases with increasing altitude, typically dropping by approximately 0.6°C for every 100-meter elevation gain ^[22]^. Lower temperatures may delay the growth cycle of tobacco plants, affecting sowing and transplanting timing. This study first investigates the relationship between altitude and agronomic traits/survival rates of tobacco plants, revealing that altitude influences effective leaf count and survival rates to some extent ^[23]^. Subsequently, a LSTM-based sowing period prediction model was developed. The model achieved 93.35 ± 2.54% accuracy across different altitude gradients, effectively predicting sowing times.

When implementing the system, transplanting dates are considered as they determine the time interval between sowing and transplanting. To ensure crops reach optimal growth stages during transplanting, sowing dates must be calculated backward from transplanting dates ^[24]^. Utilizing planned transplanting dates helps farmers schedule sowing more rationally, ensuring crops grow in the best season to enhance yield and quality. During deployment, this study reserves an API (Application Programming Interface) to enable communication and data exchange between different software systems. The API requests employ the HTTP method, ensuring broad compatibility with other platforms ^[25]^.

The results of this study show that the prediction of sowing period is very good at different altitudes, but the tobacco variety is based on the widely planted Yuntian 87, and more varieties will be considered in the future to improve the wide application of the system.

